# Efficacy of intensive seawater irrigation in mitigating climate-driven increases in incubation temperature of green sea turtle (*Chelonia mydas*) nests

**DOI:** 10.64898/2026.02.27.708658

**Authors:** David M. Adams, Sean A. Williamson, Roger G. Evans, Richard D. Reina

## Abstract

Sea turtles exhibit environmental sex determination and face risks of over-feminization, heat-induced embryonic failure, and hatchling mortality due to rising global temperatures. Mitigating these impacts of climate change may necessitate interventions to reduce sand temperature. One proposed strategy is to irrigate nests with seawater, but uncertainties exist regarding turtle egg tolerance to saline nest sand. To test the hypothesis that sea turtle embryos can tolerate a regimen of irrigation with seawater at a management-relevant scale, we investigated the impact of two levels of large-scale irrigation using cooled seawater on green turtle nests and embryos, assessing the effects on important nest environmental factors and developmental success. Irrigation which simulated 200 mm of rain reduced the temperature in clutches by up to 5.6 °C (1.34 ± 0.10 mean ± SD) without adversely affecting clutch oxygen levels, sand water potential, or sand moisture content, but our irrigation regimens resulted in very low hatching success (1.5%). However, late-stage embryonic mortality predominated, suggesting that early embryos may have an unexpected tolerance to saline sand and increasing our understanding of sea turtle resilience to seawater irrigation. The observation that younger embryos may be less susceptible to seawater-associated mortality than mature embryos near hatching further informs the limitations and potential applications of seawater irrigation as a management strategy.

## 1. Introduction

Sea turtles are a clade of marine reptiles with temperature-dependent sex determination (Yntema and Mrosovsky 1980, reviewed in Wibbels 2003) which produce a greater proportion of female hatchlings at higher incubation temperatures (Morreale et al. 1982). Contemporary climate models predict that temperatures at nesting beaches will continue to rise (Fuentes et al. 2010, IPCC 2023), posing a threat to sea turtles through increased hatchling mortality and skewed primary sex ratios (Pike 2014, Hays et al. 2017, Laloë et al. 2017). This underscores the pressing need for conservation strategies aimed at mitigating the impact of climate change on sea turtle populations, with a focus on reducing sand temperature at nesting sites.

Proposed methods to achieve this objective include shading and/or irrigation. Shading nests, either *in situ* or by translocating nests to natural shaded areas or shaded hatcheries, demands various, potentially costly investments in personnel and infrastructure (Wood et al. 2014), or risks resistance from beach users for reasons of aesthetics or public access. Watering, or ‘irrigation’, of nests, although marginally less effective at reducing temperature than shading (Hill et al. 2015), is a less labour-intensive and more cost-effective alternative to translocation (Fuentes et al. 2012). Irrigation of nests with fresh water can effectively cool sand and maintain beneficial moisture levels in nests (Jourdan and Fuentes 2015, Lolavar and Wyneken 2021). However, the limited availability of fresh water, particularly in regions where many nesting beaches are situated, poses challenges to the feasibility of this method. Seawater is abundantly available at all nesting beaches and is gaining attention as a viable alternative to fresh water for reducing sand temperature around sea turtle nests (Gatto et al. 2023).

The application of seawater can reduce nest temperature and generate male-producing conditions, if it is administered when the embryos are at the sex-determining stage (Smith et al. 2021, Young et al. 2023). However, doing so requires knowledge of where and when specific nests were laid, in order to effectively target this developmental stage. While this might be possible at small spatial and temporal scales, addressing the challenges posed by climate change to sea turtle populations may necessitate a large-scale intervention. This is particularly relevant given that managers may not have the necessary information to accurately administer water directly to nests at the most effective time. Implementing solutions at a scale necessary for effective management may involve the recurrent irrigation of expansive areas within nesting habitats (Gatto et al. 2023), but data on the effects of such a large-scale management intervention on sea turtle reproductive outcomes are lacking.

Seawater alters many important sand characteristics that affect eggs and developing embryos, including salinity, moisture, and oxygen content (Bustard and Greenham 1968, Limpus et al. 2020). Research on the effect of freshwater infiltration on nest oxygen availability has yielded mixed and sometimes conflicting results (Prange and Ackerman 1974, Booth 1998), and even less is known of the effect of seawater on nest oxygen. Extensive research has shown the detrimental effects of extreme salinity and/or moisture on turtle hatching success (Bustard and Greenham 1968, Kraemer and Bell 1980, Foley et al. 2006, Caut et al. 2010, Limpus et al. 2020), although mostly in the context of natural or simulated tidal or groundwater inundation rather than irrigation. On the other hand, Caut et al. (2010) suggested that marine turtle nests have the ability to tolerate periods of seawater inundation, provided such inundation events are not too frequent or severe. Consequently, conservation managers require a more thorough understanding of the potential impacts of irrigation with seawater on developmental success before considering the implementation of such an intervention.

At Raine Island, one of the largest green turtle rookeries in the world (Limpus and Fien 2009) and where temperatures are such that almost no males have been produced for several decades (Jensen et al. 2018, Booth et al. 2020), as much as 200 mm of simulated rainfall in a single application may be required to cool the nest environment to a male-producing temperature (Gatto et al. 2021, Gatto et al. 2023). The approach employed in previous seawater irrigation experiments by Smith et al. (2021) and Young et al. (2023) requires knowledge of the precise age and location of nests in order to irrigate them by hand during their thermosensitive period. This is unlikely to be possible at large rookeries because the exact nest location or age may not be known, or closely adjacent nests may be of different ages. In such cases, a solution would be to irrigate at a place, scale, time and frequency that has the greatest probability of cooling the greatest number of nests during their sex-determining periods to produce more males or reduce heat-induced hatchling mortality (Gatto et al. 2023).

We investigated whether using seawater to apply 200 mm of simulated ‘rainfall’ every 4 days would reduce the sand temperature around green turtle clutches by 2 to 4 °C below ambient for as long as the irrigation regimen continued. Therefore we aimed to determine the effects of a large-scale and prolonged regimen of seawater irrigation on the characteristics of the nest environment and the impact of those characteristics on embryonic development and developmental success in green sea turtles, *Chelonia mydas*. To achieve this aim, we relocated green turtle clutches to three large beach hatcheries and repeatedly irrigated them with cooled seawater as they developed, while measuring the effects of irrigation on moisture, salinity, and water potential in the sand at nest depth. We also monitored the temperature and oxygen concentration amongst the developing eggs, and determined the relative cooling effect of two different levels of irrigation, with one treatment having half the volume of water used in the other. Finally, we determined the effect of irrigation on developmental success by opening unhatched eggs and field staging the embryos.

## 2. Materials and Methods

### 2.1. Regulatory approvals

This work was approved by the Animal Ethics Committee of the School of Biological Sciences, Monash University (Approval #30901). Egg collection was conducted under the authority of a scientific research permit (P-PTUKI-100179108-1) issued by the Queensland Department of Environment and Science.

### 2.2. Study site and egg collection

We collected 2,227 *C. mydas* eggs from 26 females at Heron Island, Queensland, Australia (23.44° S, 151.91° E) from February 2–27, 2023. Eggs were relocated to beach hatcheries at Heron Island within 3.1 hours of oviposition.

### 2.3. Hatchery irrigation experiment

We assigned relocated clutches to one of three hatcheries receiving ‘intense’ irrigation (n = 721 eggs), ‘moderate’ irrigation (n = 798 eggs), or no irrigation (control, n = 708 eggs), using cooled seawater to simulate rainfall. Irrigation was conducted at intervals of 2 to 8 days (mean = 3.9). Temperature and oxygen levels within clutches were monitored using data loggers and gas sampling tubes, and sand conditions (salinity, % moisture, and water potential) were monitored using in-ground sensors (See Section 2.3.1.4). Hatching success and developmental stage at death were assessed following emergence or after at least 80 days of incubation.

#### 2.3.1. Irrigation and monitoring of nests

##### 2.3.1.1. Hatchery irrigation regimens

Relocated clutches were randomly assigned to one of three experimental hatcheries. Hatcheries were designated as ‘intense’ irrigation (simulating 200 mm of rainfall per application, n = 721 eggs from 9 clutches) or ‘moderate’ irrigation (100 mm per application, n = 798 eggs from 9 clutches). The ‘control’ hatchery plot remained unirrigated (n = 708 eggs from 8 clutches) but received natural rainfall. We assumed that the natural rainfall received by all hatcheries was the same because of their close proximity. The levels of irrigation were based upon the recommendations of Gatto et al. (2021, 2023), who suggested that as much as 200 mm of simulated rainfall may be required every 4 days to create male-producing temperatures on beaches that had experienced substantial warming over recent decades.

##### 2.3.1.2. Rainfall simulation

Rainfall was simulated using cooled seawater applied by hand-held hose with spray attachment, since the automatic irrigation devices we tested provided unreliable service when delivering salt water. Seawater was cooled to 18.2 ± 0.2 °C in a storage tank and had warmed to 19.8 ± 0.6 °C at the outflow after passing through the hose at ambient temperature. Water was pumped from the storage tanks to the hatcheries through a 25 mm hose (PVC Superflex, Dixon, Maryland, USA) and was sprayed over the sand through an adjustable diffuser nozzle (Gardena, Ulm, Germany) using a methodical back-and-forth motion. We measured the amount of water applied using a rain gauge placed in the centre of the irrigated area and ensured even coverage by placing several 500 ml plastic cups in the hatchery during irrigation and irrigating until they were filled with approximately the same amount of water.

To test the response of the sand to proposed irrigation regimen, we began pre-watering the intensely irrigated hatchery enclosure 12 days prior to relocating clutches. The purpose of the pre-watering was to determine if proposed watering schedule (200 mm about each 4 days) was sufficient to cause the desired temperature change, and if not, to adjust as necessary. The first application of water to the moderate hatchery occurred 8 days after the first clutch was relocated. This was done to reduce the risk of very early mortality (i.e., to avoid the possibility that extremely high salinity may be lethal to freshly oviposited eggs) (Limpus et al. 2020).

Results of the pilot pre-watering study and previous studies of *in situ* irrigation of nests (Hill et al. 2015, Smith et al. 2021, Young et al. 2023) indicated that it would be necessary to irrigate the hatcheries every 3–4 days to maintain temperature at nest depth at least 2 °C below natural conditions. We therefore watered, on average, every 3.9 (range 2–8) days. The target interval of 3 days was delayed on several occasions due to storms, equipment failures, or logistical conflicts. Nests were watered 14–18 times, depending on when they were placed in the hatchery. So that surface heat would not be conducted down to the clutches (Smith et al. 2021), we always allowed the surface sand to cool for at least 6 h after sunset before irrigating.

Sea turtle eggs incubated below 23 °C do not hatch (Bustard and Greenham 1968, Yntema and Mrosovsky 1980, Ackerman 1997). Consequently, irrigation continued throughout incubation until the point was reached, determined by the seasonal progression, when we had reasonable confidence that the temperature at nest depth would fall below 23 °C due to watering. At that time, irrigation was discontinued and clutches in the irrigated hatcheries were between 50.4–72.5 d old.

##### 2.3.1.3. Temperature and oxygen within clutches

Into the centre of each clutch of eggs we placed one temperature logger (HOBO MX2201, accuracy ±0.5 °C, resolution 0.04 °C, Onset Computer Corp., Bourne, MA, USA) programmed to record temperature every 15 minutes so that transient changes in temperature immediately after watering could be captured. Logger data were averaged over 24 h to calculate the average daily temperature within each clutch. To determine the cooling effect of irrigation on clutches (ΔT), we calculated the difference between the mean temperature in the hour immediately before irrigation commenced and the lowest temperature recorded before the next watering.

Into the centre of each clutch of eggs we also placed one gas sampling port made of a 40 mm hollow plastic ball perforated on the lower side and connected to 3 mm PVC tubing leading to the surface. The end of the tube was capped to prevent sand and ambient air from entering and was buried ∼5 cm under the sand when not in use. The gas sampling apparatus consisted of a 50 ml syringe equipped with a 3-way stopcock. The nest air sampling tube was connected to the inlet port of the stopcock and a pass-through oxygen sensor (PS-3217, accuracy ±1% O_2_, resolution 0.01%, PASCO Scientific, Roseville, CA, USA) was coupled to the outlet port via a sampling chamber. The air was drawn from the nest into the syringe via the sampling tube, then injected through the oxygen sensor (Supplementary Fig. 1). The stopcock ensured that air flowed only in one direction during sampling so that no surface air could enter the nest.

Gas samples were collected from each nest every second day for determination of oxygen concentration during incubation and to monitor changes in oxygen concentration that may have resulted from irrigation. First, we purged the air in the sampling tube by withdrawing and discarding ∼20 ml of air. Then, we withdrew 50 ml of nest air and injected it through the oxygen sensor. After the initial reading had stabilised (approximately 30 seconds), we tested another 50 ml of nest air. After the second sample of air, we removed the sensor from the sampling chamber, thereby exposing it to ambient air to verify calibration. Oxygen concentration was recorded at 1 s intervals throughout the sampling process and the entire process typically took 2 to 3 minutes. Prior to determining oxygen concentration for each gas sample, the oxygen sensor was calibrated to 20.9 ± 0.1% O_2_ with ambient air. We used these data to detect changes in clutch oxygen concentration that resulted from irrigation (hereafter, ‘ΔO_2_’) by comparing the oxygen level after irrigation with the level recorded before irrigation. Sampling of oxygen was scheduled so that at least 4 of the 9 irrigated nests and 3 of the 8 control nests were sampled each day, leaving the remaining nests in each hatchery to be sampled on alternate days.

##### 2.3.1.4 Hatchery sand conditions at nest depth

All hatcheries were equipped with instruments to measure sand conditions at nest depth daily. We used an in-ground sensor (TEROS-12, Meter Group, Inc., Pullman, WA, USA) to generate an index of salinity in the form of bulk electrical conductivity (EC_b_, measured in dS m^-1^, accuracy ±5% of measurement, resolution 0.001 dS m^-1^) and volumetric moisture content (accuracy ±3%, resolution 0.1%). We also installed a tensiometer (TEROS-32, accuracy ±0.15 kPa, resolution 0.0012 kPa, Meter Group, Inc.) in each hatchery to record the water potential as a quantitative measure of the tendency of water to move into or out of the eggs along the water potential gradient. One sensor of each type was placed at a depth of 60 cm in the centre of each hatchery.

The sensors used in this study to measure sand conditions at nest depth are affected by saturated conditions, rendering them unreliable in measuring both moisture content and water potential. For this reason, and so as not to collect data that reflected transient saturated sand conditions, the sand was allowed to drain for several hours after watering (mean 13.1 h, range 7.6–22.3 h) before measuring sand moisture, salinity, water potential, and temperature at nest depth.

##### 2.3.1.5. Developmental outcomes

After the hatchlings had emerged from the nests, we excavated each nest by hand to determine hatching success and embryonic stage at mortality in unhatched eggs. When nests showed no signs of emergence after at least 80 d since oviposition (mean 88.0 ± 3.3 d, mean ± SD, n = 19), we excavated and opened unhatched eggs and determined stage at mortality using an established field staging method (Leslie et al. 1996, Rafferty et al. 2011, Williamson et al. 2019) based on Miller’s (1985) 31-stage developmental chronology. Eggs that showed obvious signs of predation (e.g., by crabs or ants) were not possible to stage and were categorised as unhatched.

### 2.4. Statistical Analysis

We assessed group differences using ANOVA, Kruskal-Wallis, or Wilcoxon tests depending on data distribution, with post hoc comparisons via Tukey’s HSD or Dunn’s test with Bonferroni correction. Assumptions for parametric tests were assessed using Levene’s test for homogeneity of variance and quantile plots and normal histograms (Mangiafico 2023) for normality. All data violated the assumptions of equal variance and normality with the exception of clutch size, average daily temperature and ΔO_2_ between hatcheries (*in situ* data) and egg diameter at each sampling interval (*ex situ* data). Therefore, we used one-way analysis of variance (ANOVA) to compare differences between groups in clutch size. To include a test for maternal influence in the effect of irrigation, two-way ANOVA was used to compare average daily temperature and ΔO_2_ between and within groups, using irrigation regimen (group) and clutch (maternal identity) as independent variables. When significant differences were detected, pairwise comparisons were made using Tukey’s HSD test. Kruskal-Wallis rank sum tests were used to compare within-group differences in ΔT and clutch daily oxygen concentration and to compare between-group differences in clutch daily oxygen concentration, salinity, moisture, and water potential at nest depth as well as hatching success, stage of mortality, and percentage of depredated eggs per clutch. Differences in ΔT between the two irrigated groups were assessed using a Wilcoxon rank sum test.

Post hoc analysis of all significant Kruskal-Wallis tests were performed using Dunn’s test (Dinno 2017) with a Bonferroni correction to control the familywise error rate of multiple pairwise comparisons. All analyses were performed using R Statistical Software with RStudio (Posit Team 2023, R Core Team 2023). Two-tailed p ≤ 0.05 was considered statistically significant.

## 3. Results

### 3.1. Effects of irrigation on incubation conditions

#### 3.1.1. Clutch temperature

One temperature logger failed, so in the analysis of clutch temperature we included nine clutches from the intense irrigation hatchery and eight clutches each from the moderate irrigation and unirrigated hatcheries. We collected clutch temperature data over an average of 70.4 days from clutches that successfully hatched (n = 7) and 88.5 days from those that showed no sign of emergence (n = 18).

Average daily temperature in clutches varied significantly between the different irrigation regimens (*F* _2, 2061_ = 307.9, *p* < 0.001) (Fig. 1). Clutches in the unirrigated hatchery had the highest average daily temperature, followed by the moderate hatchery, whilst clutches in the intense hatchery were the coolest (Table 1). The control clutches were significantly warmer than both intense and moderate clutches and moderate clutches were significantly warmer than intense clutches (*p <* 0.001 in all pairwise comparisons). Within each hatchery, there was significant between-clutch variation in average daily temperature.

**Figure 1.**
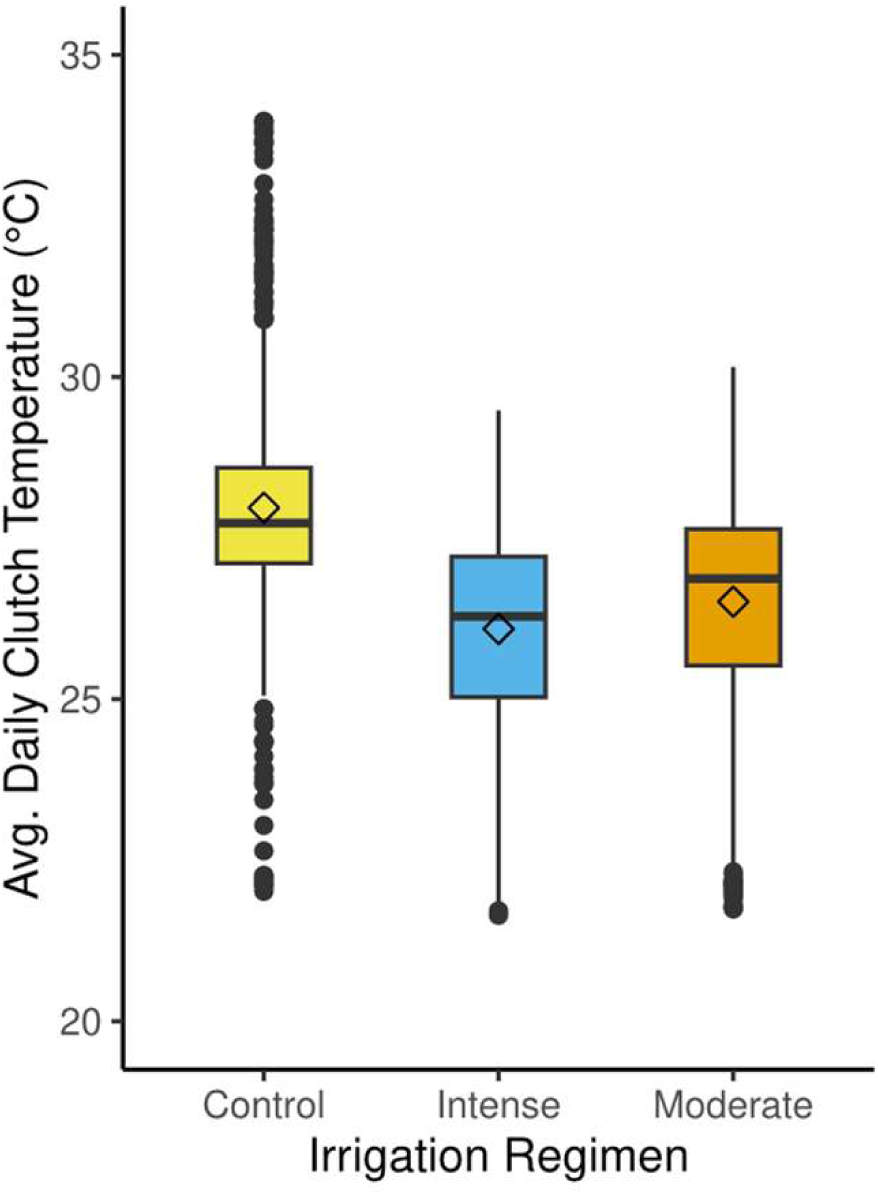
Average daily temperature of green turtle (*Chelonia mydas*) egg clutches incubating in hatcheries at Heron Island, Queensland, Australia. Nests were irrigated with 200 mm (Intense) or 100 mm (Moderate) of simulated rainfall using cooled seawater every 3.9 ± 1.3 (mean ± SD) days. Control nests were not irrigated and all nests were subject to natural rainfall. Values are the average over a 24-h period of measurements recorded every 15 minutes during incubation. All groups were significantly different according to ANOVA and post hoc Tukey’s HSD testing (α ≤ 0.05). Boxplot centre lines show the medians, diamonds are mean values, box limits indicate the 25th and 75th percentiles, whiskers extend 1.5 times the interquartile range from the 25^th^ and 75th percentiles, filled circles outside the whiskers represent outliers.

**Table 1.** Green turtle *(Chelonia mydas)* nest environmental conditions, hatchery sand characteristics, and embryonic developmental response to repeated irrigation with cooled seawater at Heron Island, Queensland. Intense irrigation simulated 200 mm of rainfall, moderate irrigation simulated I 00 mm of rainfall. Nests were watered 14 - 18 times, irrigation was applied every 3.9 ± 1.3 (mean± SD) days.

|  | Irrigation Regimen / Hatchery |  |  | Test statistic | df | *p* | Main effect size | Post hoc analysis | Clutch effects |
| --- | --- | --- | --- | --- | --- | --- | --- | --- | --- |
|  | Control | Intense | Moderate |  |  |  |  |  |  |
| Clutch temperature data |  |  |  |  |  |  |  |  |  |
| Mean # eggs per nest (# nests) | 88.5 (8) | 80.1 (9) | 88.3 (8) | $F = 0.70$ | 2, 22 | 0.51 | ns | Control = Intense = Moderate | NA |
| Clutch ADT, °C | 28.0 ± 0.08 | 26.1 ± 0.06 | 26.5 ± 0.06 | $F = 307.9$ | 2, 2061 | **< 0.001** | $\eta^2_{\text{partial}} = 0.23$ (m) | Control > Moderate > Intense | **Yes** |
| Clutch ΔT post-irrigation, °C | NA | -1.34 ± 0.10 | -0.84 ± 0.08 | $W = 6459$ | 1 | **< 0.001** | $r = 0.30$ (m) | Intense > Moderate | No |
| Clutch oxygen data |  |  |  |  |  |  |  |  |  |
| Mean # eggs per nest (# nests) | 88.5 (8) | 80.1 (9) | 88.7 (9) | $F = 0.78$ | 2, 23 | 0.47 | ns | Control = Intense = Moderate | NA |
| Oxygen concentration, % | 19.9 ± 0.05 | 20.3 ± 0.02 | 20.2 ± 0.02 | $\chi^2 = 16.41$ | 2 | **< 0.001** | $\eta^2 = 0.01$ (s) | Control < Intense = Moderate | **Yes** |
| ΔO<sub>2</sub>, % | NA | -0.03 ± 0.01 | -0.04 ± 0.02 | $F = 0.09$ | 1, 286 | 0.77 | ns | Intense = Moderate | No |
| Hatchery sand at nest depth |  |  |  |  |  |  |  |  |  |
| Salinity (EC<sub>b</sub>), dS m<sup>-1</sup> | 0.03 ± 0.03 | 0.543 ± 0.092 | 0.501 ± 0.215 | $\chi^2 = 177.9$ | 2 | **< 0.001** | $\eta^2 = 0.56$ (L) | Control < Intense = Moderate | NA |
| Moisture, % (v/v) | 10.2 ± 1.12 | 13.8 ± 1.58 | 13.8 ± 2.11 | $\chi^2 = 173.9$ | 2 | **< 0.001** | $\eta^2 = 0.55$ (L) | Control < Intense = Moderate | NA |
| Water potential, kPa | -7.30 ± 4.01 | -3.16 ± 0.37 | -2.61 ± 0.38 | $\chi^2 = 172.9$ | 2 | **< 0.001** | $\eta^2 = 0.55$ (L) | Control < Moderate < Intense | NA |
| Developmental success |  |  |  |  |  |  |  |  |  |
| Hatching success, % | 72.3 ± 12.9 | 1.79 ± 0.92 | 1.18 ± 0.72 | $\chi^2 = 15.1$ | 2 | **< 0.001** | $\eta^2 = 0.57$ (L) | Control > Intense = Moderate | NA |
| Mortality at field stage 0, % | 46.0 ± 14.9 | 30.4 ± 5.5 | 28.5 ± 5.4 | $\chi^2 = 0.469$ | 2 | 0.79 | ns | Control = Intense = Moderate | NA |
| Mortality at field stage 1, % | 1.0 ± 0.8 | 1.7 ± 0.5 | 4.5 ± 1.5 | $\chi^2 = 8.09$ | 2 | **0.02** | $\eta^2 = 0.26$ (L) | Control = Intense < Moderate | NA |
| Mortality at field stage 2, % | 0 ± 0 | 4.4 ± 1.0 | 5.5 ± 1.5 | $\chi^2 = 13.4$ | 2 | **< 0.01** | $\eta^2 = 0.50$ (L) | Control < Intense = Moderate | NA |
| Mortality at field stage 3, % | 40.5 ± 14.7 | 63.6 ± 5.2 | 61.5 ± 6.3 | $\chi^2 = 1.12$ | 2 | 0.57 | ns | Control = Intense = Moderate | NA |
| Depredated eggs per clutch, % | 4.5 ± 1.5 | 3.2 ± 1.8 | 9.7 ± 3.6 | $\chi^2 = 2.90$ | 2 | 0.23 | ns | Control = Intense = Moderate | NA |

Data presented are clutch average daily temperature (ADT), change in clutch temperature after irrigation ( $\Delta T$ ), clutch daily oxygen concentration, change in clutch daily oxygen after irrigation ( $\Delta O_2$ ), hatchery sand conditions at nest depth (60 cm), and developmental success of hatchery-incubated clutches. Data on clutch ADT and clutch $\Delta T$ were calculated using one less clutch in the moderate hatchery than on % oxygen, $\Delta O_2$ , and developmental success, so sample sizes are given at the beginning of the relevant sections. Data are presented as means ± SE. Test statistics reported are the results of ANOVA ( $F$ ), Kruskal-Wallis rank sum tests ( $\chi^2$ ), and Wilcoxon rank sum tests ( $W$ ). Effect sizes are reported as small (s), moderate (m), or large (L) unless no significant differences were detected (ns). Post hoc pairwise comparisons were made using Tukey's HSD (ANOVA) or Dunn's test with Bonferroni-corrected $p$ -values (Kruskal-Wallis). NA = not applicable.

Thee cooling effect of irrigation on clutches (ΔT, defined as the departure from pre-watering temperature) in all clutches ranged from -5.58 °C to +0.02 °C and was significantly different between the two irrigated hatcheries (*W* = 6459, *p* < 0.001) (Fig. 2). Mean ΔT in the intense hatchery was greater than in the moderate hatchery (Table 1). There was no significant difference in ΔT between clutches in the same hatchery.

**Figure 2.**
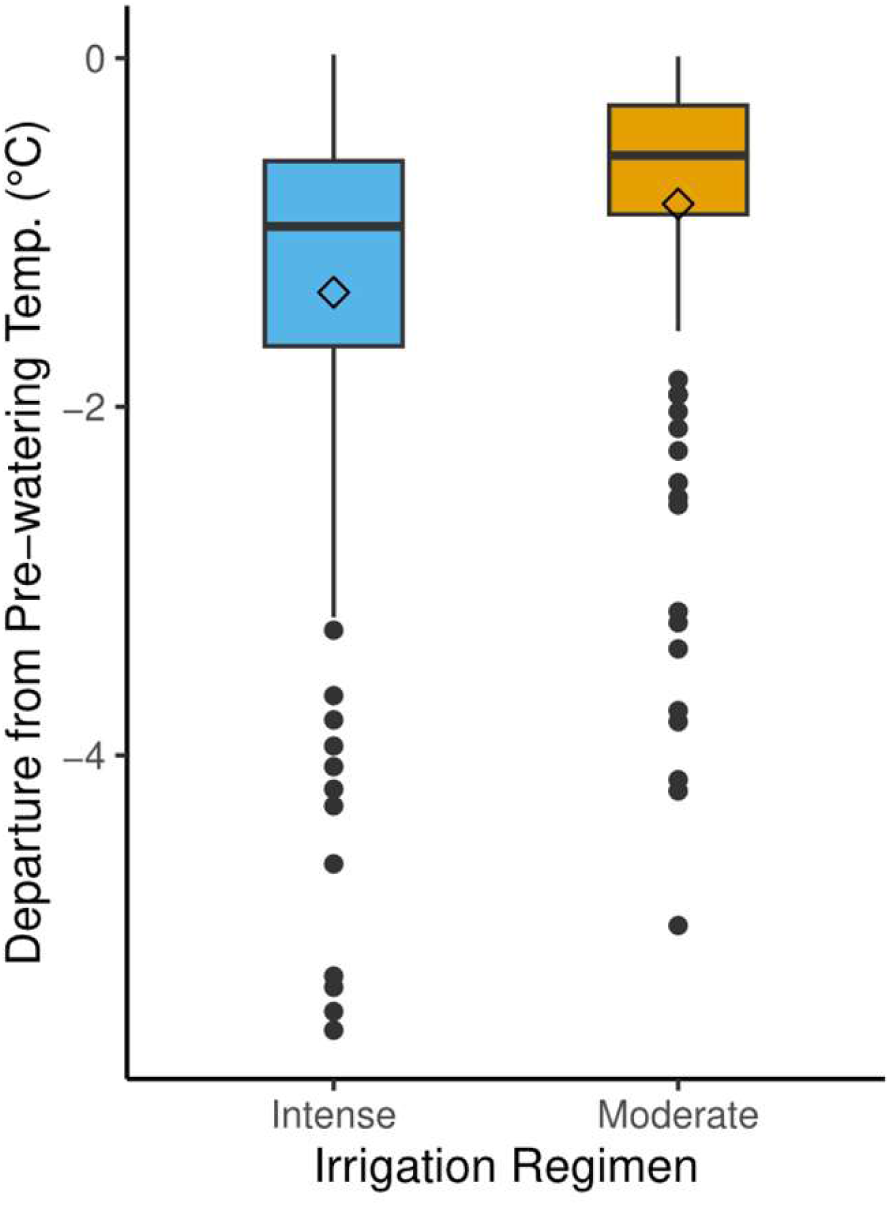
Cooling effect of seawater irrigation on green turtle (*Chelonia mydas*) egg clutches at Heron Island, Queensland, Australia. Nests were irrigated with 200 mm (Intense) or 100 mm (Moderate) of simulated rainfall using cooled seawater every 3.9 ± 1.3 (mean ± SD) days. Nests were subject to natural rainfall. Values are the difference between the lowest temperature recorded after irrigation and the average temperature in the hour before irrigation. The two groups were significantly different according to a Wilcoxon rank sum test (*p* < 0.001). Boxplot centre lines show the medians, diamonds are mean values, box limits indicate the 25th and 75th percentiles, whiskers extend 1.5 times the interquartile range from the 25^th^ and 75th percentiles, filled circles outside the whiskers represent outliers.

#### 3.1.2. Clutch oxygen concentration

The patterns of O_2_ concentration during incubation varied with irrigation regimen. During the first half of incubation, clutch O_2_ in all hatcheries was similar, but in the second half of incubation control clutches had lower O_2_ than clutches in either of the irrigated hatcheries (Fig. 3). Oxygen concentration was lowest in the control hatchery and varied significantly between irrigated hatcheries and control (χ^2^ = 16.41, df = 2, *p* < 0.001), but not between the two irrigated hatcheries (*p* = 0.80) (Table 1, Fig. 4). Within each hatchery, oxygen concentration varied significantly between clutches (Supplementary Table 1). Irrigation resulted in a ΔO_2_ value that ranged from -0.5% to +0.6% in intense clutches and from -0.8 to +0.6 in moderate clutches (Fig. 5), but were not statistically significant (*F* _1, 286_ = 0.296, *p* = 0.77). There were no significant differences in ΔO_2_ values between clutches in the same hatchery.

**Figure 3.**
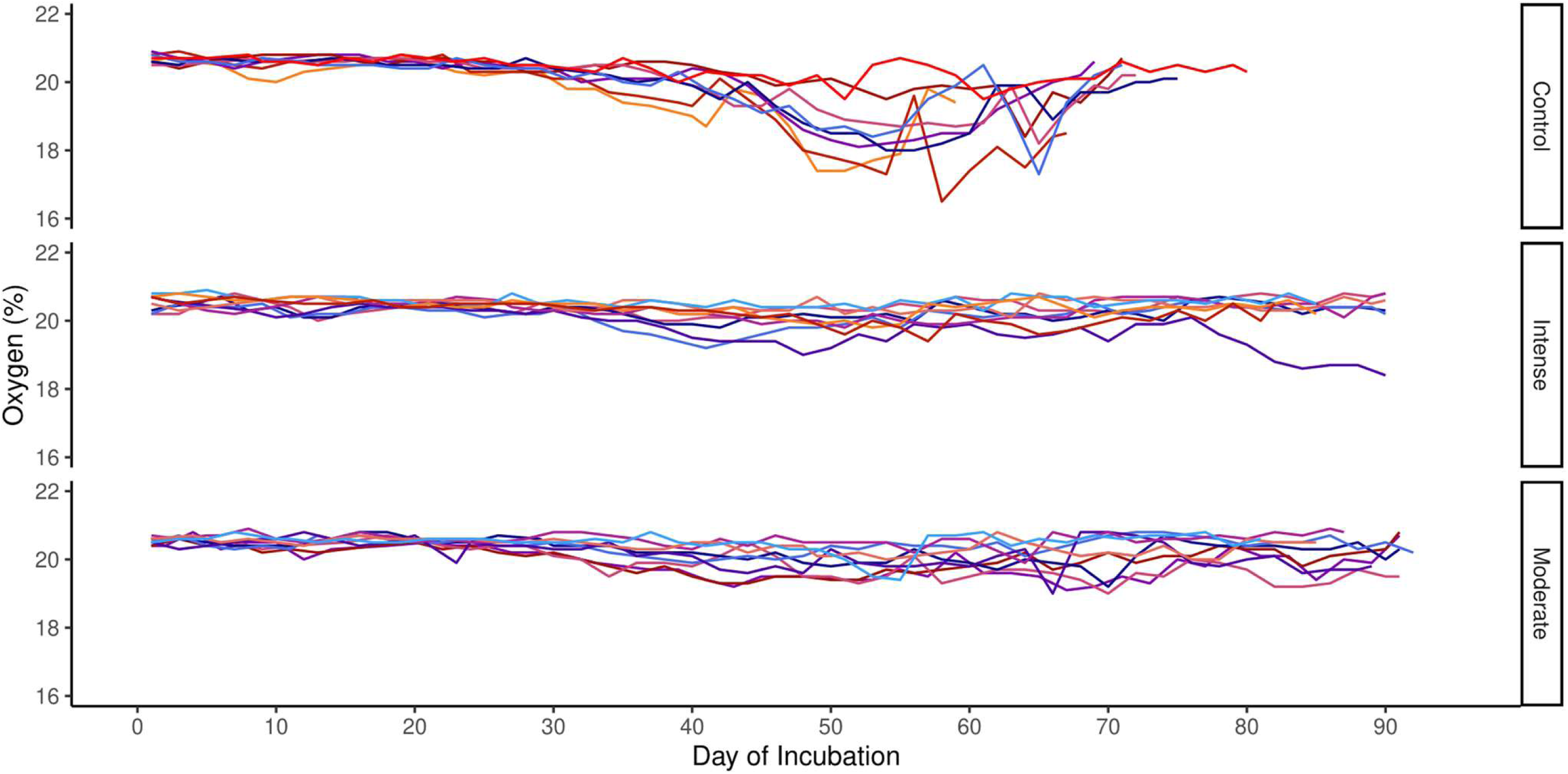
Oxygen concentration (%) in clutches of green turtle (*Chelonia mydas*) eggs during incubation at Heron Island, Queensland, Australia. Irrigation regimens simulated 200 mm (intense) or 100 mm (moderate) of rainfall. Nests were irrigated with cooled seawater every 3.9 ± 1.3 (mean ± SD) days. Control nests were not irrigated and all nests were subject to natural rainfall. Oxygen concentration was recorded once every 2 days during incubation. Patterns of oxygen consumption were similar for the first half of incubation, but high mortality in the irrigated nests and high hatching success in the control nests resulted in significant differences in oxygen concentration between the irrigated groups and the control group (Kruskal-Wallis and Dunn’s post hoc test, *p* < 0.001).

**Figure 4.**
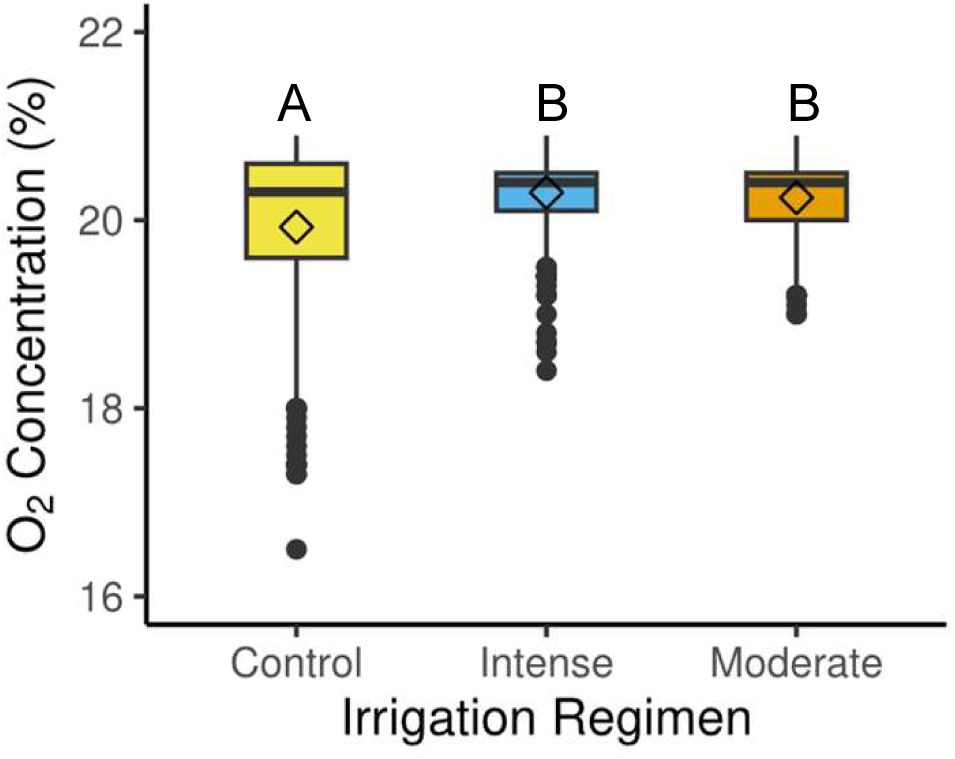
Daily oxygen concentration (%) in clutches of green turtle (*Chelonia mydas*) eggs during incubation in hatcheries at Heron Island, Queensland, Australia. Irrigation regimens simulated 200 mm (intense) or 100 mm (moderate) of rainfall. Nests were irrigated with cooled seawater every 3.9 ± 1.3 (mean ± SD) days. Control nests were not irrigated and all nests were subject to natural rainfall. Oxygen concentration was recorded once every 2 days during incubation. Letters above boxplots indicate statistical groups; when letters are the same, oxygen concentration did not significantly differ between irrigation regimens according to Kruskal-Wallis and Dunn’s post hoc test (α ≤ 0.05). Boxplot centre lines show the medians, diamonds are mean values, box limits indicate the 25th and 75th percentiles, whiskers extend 1.5 times the interquartile range from the 25th and 75th percentiles, filled circles outside the whiskers represent outliers.

**Figure 5.**
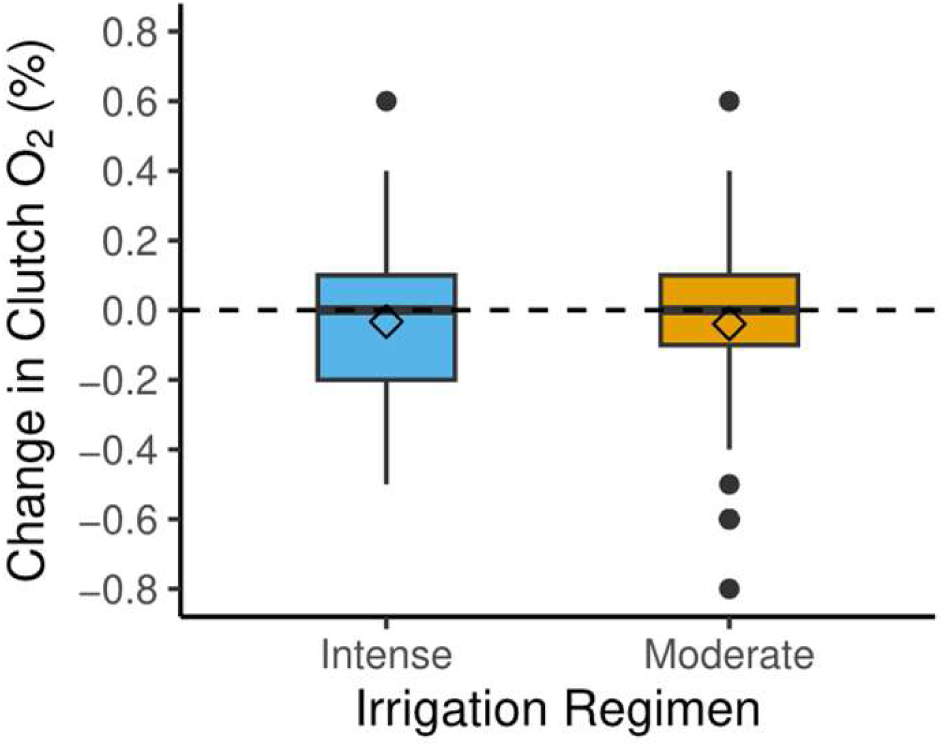
Changes in percent oxygen concentration as a result of irrigation with cooled seawater (ΔO2) in clutches of green turtle (*Chelonia mydas*) eggs during incubation in hatcheries at Heron Island, Queensland, Australia. Irrigation regimens simulated 200 mm (intense) or 100 mm (moderate) of rainfall. Nests were irrigated with cooled seawater every 3.9 ± 1.3 (mean ± SD) days and all nests were subject to natural rainfall. Oxygen concentration was recorded once every 2 days during incubation. Values shown are difference between the lowest oxygen level measured after irrigation and the level recorded before irrigation. There was no significant difference between irrigated groups (ANOVA, p = 0.77). Dashed horizontal line indicates zero net change in oxygen concentration. Boxplot centre lines show the medians, diamonds are mean values, box limits indicate the 25th and 75th percentiles, whiskers extend 1.5 times the interquartile range from the 25th and 75th percentiles, filled circles outside the whiskers represent outliers

#### 3.1.3. Salinity

Prior to irrigation, salinity (EC_b_) at nest depth in all hatcheries was 0.004–0.019 dS m^-1^. Salinity in the control (unirrigated) hatchery ranged from 0.004–0.141 dS m^-1^, with the most saline conditions recorded after substantial natural rainfall (>50 mm). During incubation of relocated clutches, sand salinity in the intense hatchery ranged from 0.180–0.855 dS m^-1^ and from 0.004–0.902 dS m^-1^ in the moderate hatchery (Fig. 6a). These differences were significant (χ^2^ = 177.9, df = 2, *p* < 0.001), but only between the control hatchery and the two irrigated hatcheries (both *p* < 0.001), not between the intense and moderate hatcheries (*p* > 0.99). The salinity in irrigated hatcheries showed a transient, short-term rise in response to irrigation with seawater, followed by a sharp decline to pre-irrigation levels in ∼3 days (Fig. 7a).

**Figure 6.**
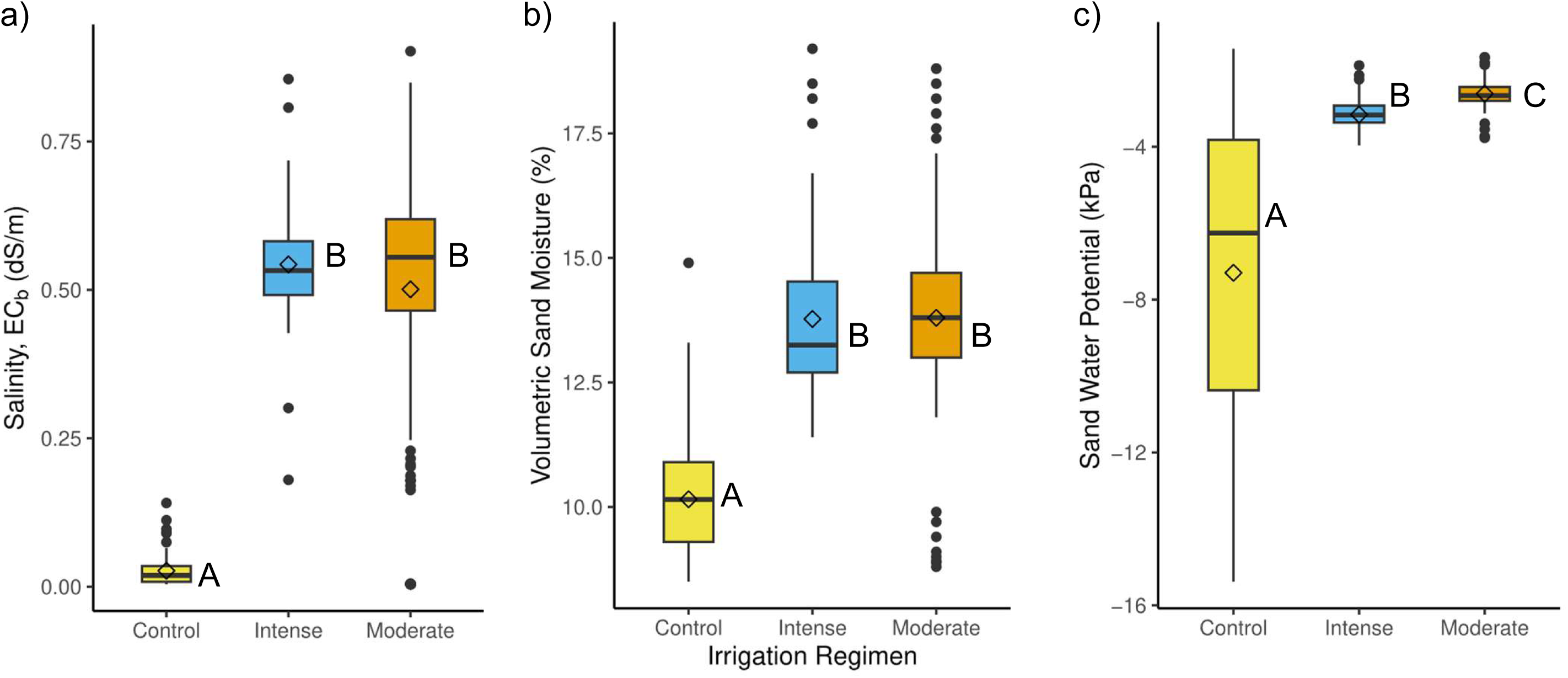
Variation in characteristics of hatchery sand at nest depth (60 cm) as a result of multiple bouts of irrigation with cooled seawater at Heron Island, Queensland, Australia. Irrigation regimens simulated 200 mm (intense) or 100 mm (moderate) of rainfall. Nests were irrigated with cooled seawater every 3.9 ± 1.3 (mean ± SD) days. Control nests were not irrigated and all nests were subject to natural rainfall. Shown here are a) sand salinity, measured as bulk electrical conductivity (ECb, see text), b) sand percent water content by volume, and c) water potential. Letters beside boxplots indicate statistical groups; when letters are the same the measured variables did not significantly differ between irrigation regimens according to Kruskal-Wallis and Dunn’s post hoc test (α ≤ 0.05). Boxplot centre lines show the medians, diamonds are mean values, box limits indicate the 25th and 75th percentiles, whiskers extend 1.5 times the interquartile range from the 25th and 75th percentiles, filled circles outside the whiskers represent outliers.

**Figure 7.**
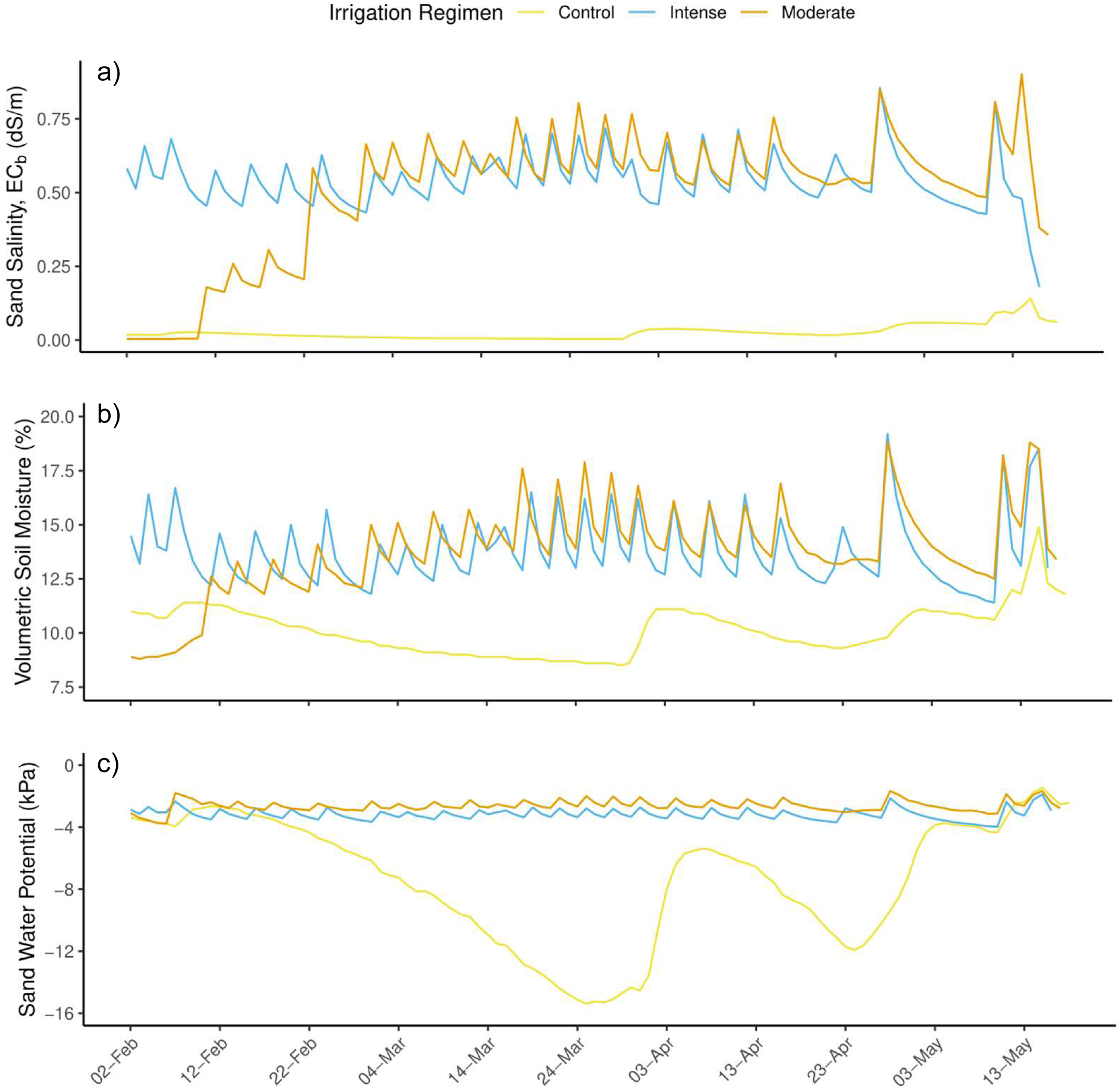
Characteristics of hatchery sand at nest depth (60 cm) during incubation of green urtle (*Chelonia mydas*) clutches under three different seawater irrigation regimens at Heron Island, Queensland, Australia. Shown here are daily measurements of a) sand salinity, measured as bulk electrical conductivity (ECb, see text), b) sand percent moisture content by volume, and c) sand water potential (matric potential plus pressure potential). Irrigation regimens simulated 200 mm (intense) or 100 mm (moderate) of rainfall. Control nests were not irrigated and all nests were subject to natural rainfall. Hatcheries were irrigated with cooled seawater every 3.9 ± 1.3 (mean ± SD) days until 16 April, beginning on 22 Jan (intense) or 11 Feb (moderate). Bouts of irrigation can be clearly seen as transient changes in all measured variables in the intense and moderate hatchery traces. Hatcheries were irrigated 1 d apart until 22 March, after which irrigation occurred on the same day. Substantial rainfall events (> 15 mm) occurred on 30 March, 28 April, 10-11 May, and 14-15 May.

#### 3.1.4. Moisture content

Hatchery sand moisture at nest depth showed a pattern similar to that of sand salinity, that is, a sharp and transient rise in moisture immediately after watering, followed by a decrease to pre-watering levels in 2 to 3 days (Fig. 7b). Sand moisture in the intense hatchery remained between 11.4%–19.2% (v/v) and the moderate hatchery ranged between 8.8%–18.8% (v/v) (Fig. 6b). There was a significant difference in sand moisture (χ^2^ = 173.9 df = 2, *p* < 0.001) between the control hatchery and both irrigated hatcheries (both *p* < 0.001), but not between the intense and moderate hatcheries (*p* = 0.99).

#### 3.1.5. Water potential

Water potential varied significantly between different irrigation groups (χ^2^ = 172.9, df = 2, *p* < 0.001) and followed a pattern in which both irrigated hatcheries had very stable conditions but the control hatchery was highly variable (Fig. 7c). Mean water potential was -3.16 kPa (range -3.97 to -1.87 kPa) in the intense hatchery and -2.61 kPa (range -3.77 to -1.66 kPa) in the moderate hatchery (Fig. 6c, Table 1). The control hatchery was subject to natural drying between rain events and thus had a wide range of water potential from -15.4 to -1.44 kPa (Fig. 6c).

### 3.2. Effect of irrigation on embryonic development

#### 3.2.1. Hatching success

Irrigation with seawater significantly reduced hatching success. In the unirrigated hatchery, 72.3% of eggs hatched, whilst only 1.8% of eggs hatched after the intense irrigation regimen and 1.2% of eggs after moderate irrigation (Fig. 8a). The difference in hatching success was significant (χ^2^ = 15.1, df = 2, *p* < 0.001) between the intense and control hatcheries (*p* = 0.004) and the moderate and control hatcheries (*p* = 0.001) but not between the intense and moderate hatcheries (*p* = 0.99). Hatching success of unirrigated clutches in the control hatchery was 82.5% when excluding the single clutch with complete failure, as it was the only clutch to be inundated by seawater multiple times due to storm activity.

**Figure 8.**
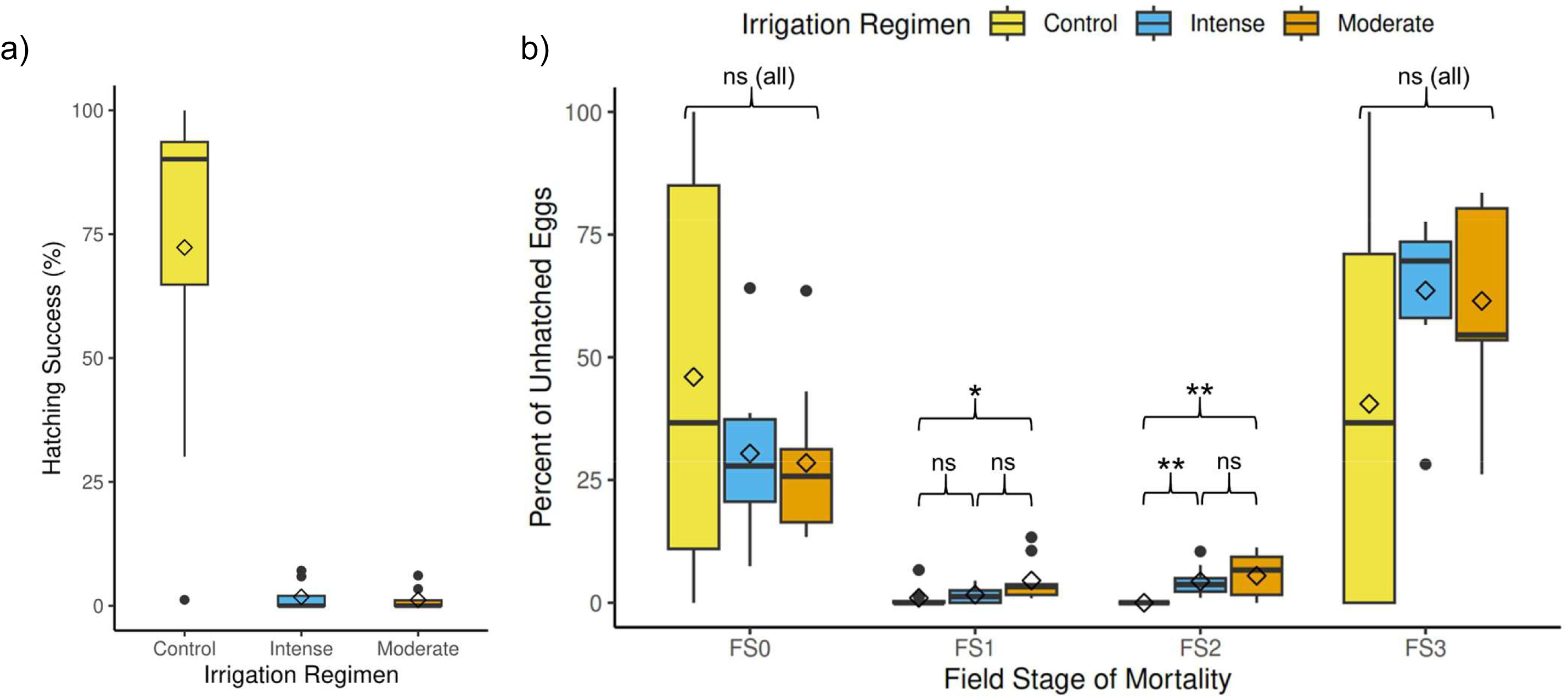
Hatching success (a) and stage of mortality of unhatched eggs (b) incubated in green turtle (*Chelonia mydas*) hatcheries irrigated with seawater at Heron Island, Queensland, Australia. Irrigation regimens simulated 200 mm (intense) or 100 mm (moderate) of rainfall. Hatcheries were irrigated with cooled seawater every 3.9 ± 1.3 (mean ± SD) days. Control nests were not irrigated and all nests were subject to natural rainfall. Hatching success (a) was significantly higher in control nests than in irrigated nests (p < 0.001) but there was no significant difference between nests under intense and moderate irrigation (p = 0.99). Significant differences in stage of mortality (b) at field stages 1 (FS1) and 2 (FS2) are indicated by asterisks (* = p ≤ 0.05, ** = p ≤ 0.01, ns = not significant). All analyses were performed with Kruskal-Wallis and post hoc Dunn’s tests, α = 0.05. Boxplot centre lines show the medians, diamonds are mean values, box limits indicate the 25th and 75th percentiles, whiskers extend 1.5 times the interquartile range from the 25th and 75th percentiles, filled circles outside the whiskers represent outliers. Note the single control nest that was repeatedly inundated by storm-related tides as an extreme outlier in (a).

#### 3.2.2. Stage of mortality

We report the stage of mortality as the proportion of unhatched eggs that died at each stage. Live and dead embryos that had pierced the eggshell but had not escaped from the egg (‘pipped’ eggs) were categorised as field stage 3, and depredated eggs were categorised as unhatched but excluded from comparisons of stage of mortality. Failed eggs in all hatcheries showed a consistent pattern in which early and late stage embryo mortality predominated and mortality at intermediate stages was rare (Fig. 8b). Over 60% of unhatched eggs in both of the irrigated hatcheries died at field stage 3. There were significant differences in the percentage of unhatched eggs that failed at stage 2 between the control hatchery and both the intense (*p* = 0.004) and moderate (*p* = 0.001) hatcheries, as well as in the percentage of embryos that died at stage 1 between the clutches in the moderate irrigation regimen and the control hatchery (*p* = 0.01) (Table 1). There were no significant between-group differences in mortality at stages 0 and 3.

## 4. Discussion

Increasing global temperature is expected to be the most important factor affecting sea turtle rookeries in the long term (Fuentes et al. 2011), underscoring the importance of management interventions directed at mitigating elevated nest temperatures. Additionally, it highlights the need for a better understanding of the consequences arising from these interventions on sea turtle populations. The results of our investigation show that irrigation with seawater can reduce the temperature of sea turtle nests, because irrigated nests were significantly cooler than unirrigated nests and the magnitude of cooling increased with the volume of water applied. Further, the amount of oxygen available to the eggs and embryos was not affected by irrigation. However, an irrigation regimen consisting of multiple applications of seawater of the magnitude employed in this study significantly altered the nest environment and had a strong negative impact on developmental success. Compared with unirrigated sand, sand in irrigated hatcheries had significantly higher salinity, moisture, and water potential. The negative impact of high salinity on early embryonic development was also evidenced by the strong reduction in hatching success that resulted from irrigation with seawater, whereby only 1.5% of all eggs in the irrigated hatcheries produced live, emergent hatchlings. These results indicate that large-scale and long-term irrigation with seawater, when applied in a regime similar to that in this study, is unlikely to be a viable solution for mitigating the effects of a warming climate on sea turtle reproductive outcomes, at least in the absence of concomitant mitigating actions. Nevertheless, our results indicate that as part of a suite of interventions, seawater irrigation may have a valuable role in reducing the effects of a warming nest environment if the intensity, frequency, timing and duration of irrigation is carefully planned and managed. We think that it is therefore worthy of further investigation and refinement in an effort to achieve this management goal.

### 4.1. Abiotic factors of the nest environment in response to seawater irrigation

#### 4.1.1. Temperature

The ability of water flux to reduce the temperature of sea turtle nests has been extensively explored, both as a result of rainfall (Houghton et al. 2007, Lolavar and Wyneken 2015, Staines et al. 2020, Laloë et al. 2021) and artificial irrigation (Hill et al. 2015, Jourdan and Fuentes 2015, Lolavar and Wyneken 2021, Smith et al. 2021, Gatto et al. 2023, Young et al. 2023). Our research builds upon that knowledge and provides further evidence that irrigation with seawater can be used to reduce the temperature of sea turtle nests. Our findings indicate that an intense level of simulated rainfall (200 mm) is more effective at cooling than a moderate level (100 mm), but that both are capable of achieving a 2 to 4 °C reduction in nest temperature (ΔT) under certain conditions. This is an encouraging result and suggests that irrigation with seawater may yet have a role to play in reducing embryonic mortality and increasing the production of male hatchlings. Further research is required to determine the appropriate frequency and volume of seawater to apply to ensure acceptable hatching success.

As expected, the average daily temperature of unirrigated clutches was significantly warmer than irrigated nests, even though at our study location the control hatchery was perforce situated high on the beach in close proximity to a stand of large coastal she-oaks (*Casuarina equisetifolia*) and was thus more shaded than the irrigated hatcheries. Given the poor hatching success in the irrigated hatcheries and the high hatching success in the unirrigated hatchery, it is likely that the temperature difference between hatcheries is due not only to irrigation but also to metabolic heating in unirrigated clutches in the later stages of incubation (Morreale et al. 1982, Broderick et al. 2001). It is also interesting to note that the mean temperature of 28.0 °C in the control clutches suggests a strong likelihood that many of the hatchlings were male, as the pivotal range of temperatures lies between 27.6 °C and 29.0 °C at this location (Ackerman 1997, Limpus and Fien 2009).

#### 4.1.2. Oxygen

We observed two distinct aspects of oxygen concentration in green turtle clutches. The main focus was on assessing the impact of irrigation on oxygen availability by determining ΔO_2_ following irrigation. However, through data collection every second day, we concurrently achieved the objective of observing the pattern of oxygen concentration throughout the entire incubation period. ΔO_2_ data revealed that neither the intense nor the moderate irrigation regimen significantly altered the oxygen concentration in the centre of the clutches. Sufficient oxygen availability is vital for successful embryonic development and the finding that irrigation does not significantly affect oxygen availability in clutches suggests that some form of irrigation may be a viable management intervention if properly executed. This conclusion is supported by numerous reports of good hatching success after relatively small-scale irrigation with both fresh water and seawater (Hill et al. 2015, Jourdan and Fuentes 2015, Lolavar and Wyneken 2021, Smith et al. 2021, Young et al. 2023).

Our observation that clutch oxygen concentration was unaffected by irrigation differs from previous research. For example, Prange and Ackerman (1974) observed a transient yet distinct rise in oxygen in a *C. mydas* nest on the Caribbean coast of Costa Rica following a heavy rain event. In contrast, Booth (1998), in his study of broad-shelled freshwater turtles *Chelodina expansa*, noted a nest becoming moderately hypoxic after a heavy rain shower. Similarly, observations on a freshwater crocodile (*Crocodylus johnstoni*) nest by Whitehead (1987) revealed a brief period of hypoxia following intense rainfall. These disparities could stem from variations in nest depth, soil type, soil organic content, and drainage characteristics of these species’ nesting habitat, but they may also be influenced by the temporal resolution of the oxygen concentration data. Prange & Ackerman (1974) collected data continuously and Booth’s (1998) sampling interval was implied to be daily, although it was not specifically identified. We measured the oxygen level in each clutch every second day due to the number of clutches being monitored and the limitation in available human resources. As a result, the time elapsed between irrigation and measurements of clutch oxygen concentration was 8–40 hours. At that temporal resolution, we did not detect a significant effect of irrigation on clutch oxygen. Nevertheless, Prange & Ackerman (1974) and Booth (1998) concluded that the observed change in available oxygen had no effect on embryonic survival and, at most, may have slowed embryonic growth during brief periods of hypoxia. We cannot exclude the possibility that clutch oxygen may have been affected for periods that were too brief to be detected by our method. Even so, considering the evidence of the current study and previous data, it seems reasonable to conclude that oxygen availability was not a limiting factor in the development of the irrigated embryos.

Under normal conditions, oxygen concentration during incubation of green turtle clutches typically follows a pattern in which O_2_ remains at or near atmospheric sea-level conditions (>19% O_2_) for the first half of incubation, after which it slowly declines to its lowest level (approximately 12–19%) in the days before hatching (Prange and Ackerman 1974; this study) (Supplementary Fig. 2). We observed this pattern in control hatchery clutches but not in the intensely irrigated clutches and only weakly in the moderate clutches. Interestingly, the lowest level of oxygen we observed in irrigated clutches, and the highest minimum oxygen of the control clutches, was 18.4%. We recorded this in the clutch that had the highest hatching success of the moderate irrigated nests (7.1%) and the lowest hatching success of the control nests (30.1%), excluding the failed control clutch that was repeatedly inundated by high tides. The observed parallels in the temporal oxygen patterns among hatcheries, coupled with the significant proportion of embryos experiencing mortality at field stage 3 of development, indicate that embryonic development in some of the irrigated clutches proceeded for a minimum of 30 days and in some cases much longer. If repeatable, making adjustments to the irrigation regimen(s) employed in this study (e.g., fewer bouts of irrigation and/or the occasional use of fresh water to flush salts from the nest environment) may facilitate greater embryonic survival beyond field stage 3, establishing irrigation with seawater as a promising strategy for sea turtle management.

#### 4.1.3. Salinity

For sea turtles, sand salinity is an important factor influencing both nest site selection and reproductive success (Bustard and Greenham 1968, Mortimer 1990, Foley et al. 2006, Limpus and Fien 2009). We irrigated the surface sand in two green turtle hatcheries with a large quantity of seawater, which is akin to simulating a minor tidal inundation event. With each application of seawater, we observed a sharp rise in the salinity of the sand, which quickly declined but nevertheless remained higher than the pre-irrigation level. The elevated salinity persisted until the sand had been flushed with substantial fresh water (i.e., natural rainfall), at which time it returned to the level observed before irrigation began. Sea turtle clutches are sensitive to tidal inundation, particularly very early or very late in development (Limpus et al. 2020), and generally suffer complete mortality if inundations persist for an extended period of time (e.g., persistent rainstorms, Bustard 1972, Dean and Talbert 1975) or over multiple occasions (Leslie et al. 1996, Caut et al. 2010, Sari and Kaska 2017). In the case of very early embryos, mortality under these circumstances results from the elevated salinity disturbing the osmotic balance between eggs and sand, leading to desiccation and mortality (Limpus et al. 2020). Although there are little data concerning the upper limit of sea turtle embryonic tolerance for salinity, the precautionary principle advises managers to maintain hatchery sand salinity as close to its baseline value as possible.

Bulk electrical conductivity (EC_b_), which can be a useful proxy for sand salinity (Valdés et al. 2014), was observed to be 0.004–0.019 dS m^-1^ in the natural beach sand at Heron Island. EC_b_ never exceeded 0.14 dS m^-1^ in the control hatchery, which, excluding the nest that was inundated by storm-driven tides, saw 82.5% hatching success. This level of EC_b_ can therefore be assumed to support good hatching success.

Rainfall affects the level of salinity in beach sand, even when the sand is above the high tide line. For example, EC_b_ in the control (unirrigated) hatchery was observed to rise from 0.02 to 0.14 dS m^-1^ after a series of substantial rainfall events. We observed a similar pattern of transient increases in sand salinity after rainfall events during range-finding experiments conducted at Heron Island in 2022 (Adams, unpublished data). Also during that pilot study, one extreme rainfall event (>250 mm) restored sand that had been irrigated with seawater (∼0.4 dS m^-1^) to pre-watering conditions (<0.05 dS m^-1^) overnight. In the current study, the general trend was for the salinity of the sand in both irrigated hatcheries to return to its pre-watering level before we irrigated again. These patterns suggest that the well-drained sand of the nesting beach facilitates a naturally occurring variation in sand salinity in sea turtle nests. The most likely explanation is that sea spray accumulates on the surface sand between rain events and percolates down through the sand column after rainfall. This phenomenon causes EC_b_ in natural nests at Heron Island to rise as high as 0.14 dS m^-1^, and perhaps higher after extended periods without rain, with no negative impacts on hatching success.

#### 4.1.4. Moisture

Moisture, water potential, and salinity are closely linked parameters of the nesting environment which substantially impact incubating eggs (Bustard and Greenham 1968, McGehee 1990, Ackerman 1997, Lolavar and Wyneken 2017). High moisture in turtle nests is associated with larger and heavier hatchlings and increased locomotor performance compared with hatchlings incubated in drier conditions (reviewed in Gatto and Reina 2022). However, excess moisture can negatively impact hatching success and even lead to complete clutch failure (Ragotzkie 1959, Caut et al. 2010, Limpus et al. 2020). Our data indicate that, at the average depth of green turtle nests, the biogenic sand of Heron Island stabilises at approximately 10% moisture (v/v). That is, the average moisture of the sand in the unirrigated hatchery was 10.2% and remained between 8.5–12.0% except for a brief period after a substantial rain event. As indicated by the high hatching success of control clutches, this level of moisture can be assumed to be within an optimal range for green turtles at Heron Island. However, it is reasonable to assume that eggs and embryos may be resistant to a greater level of moisture, such as was seen in the irrigated hatcheries, for limited periods (McGehee 1990, Limpus et al. 2020).

In the irrigated hatcheries, the levels of irrigation applied are presumed to have nearly saturated the sand during watering, but the highly permeable sand drained rapidly. The data reveal that soil moisture returned to ∼15.5% within one day of irrigation and subsequently declined by up to 2% per day. Further evidence of this is provided by opportunistic measurements of hatchery sand moisture taken immediately after irrigation. On one occasion we observed a maximum post-irrigation soil moisture of over 37%, which fell to 25% one hour after irrigation and was 13.8% 12 hours later (data not shown). On another occasion, moisture rose from 12.9% immediately before irrigation to 49.4% immediately after and declined to 15.0% within 12 h. From this we can infer that the sand at Heron Island has a high hydraulic conductivity which maintains soil moisture below ∼15% except during periods of high water flux, such as during heavy rain or tidal inundation.

Buried reptile eggs are generally tolerant to soil moisture variation and can develop successfully over a wide range of moisture content (McGehee 1990, Ackerman and Lott 2004). For example, McGehee (1990) found that hatching success of loggerhead sea turtle (*Caretta caretta*) eggs incubated in sand at a constant 12% moisture (w/w) was not significantly different to eggs in dry sand (0% moisture) or eggs in sand at 6% moisture. However, they found hatching success to be significantly reduced at 18% and 24% moisture and concluded that embryos may be more tolerant of low moisture than very high moisture. The mean moisture in both irrigated hatcheries was 13.8%, but moisture in the irrigated hatcheries exceeded that level for brief periods after irrigation. This may have contributed to the low hatching success we observed. Research into the upper limit of moisture tolerance would provide important data on reptile physiology and inform management decisions regarding the intensity and duration of any proposed irrigation program.

#### 4.1.5. Water potential

The water balance of incubating eggs is the result of primarily abiotic forces which facilitate the movement of water down concentration gradients in both liquid and vapor phases (Packard and Packard 1988). Barring any impermeable barriers, water moves from areas of high water potential (wet areas) to areas of lower water potential (dry areas). Total water potential in a soil matrix (in this case, beach sand) is the sum of matric, osmotic, pressure, gravitational, and overburden potentials (Papendick and Campbell 1981). Matric potential (adsorption between water and soil particles, i.e., surface tension) and osmotic (solute) potential are the largest components, whilst pressure potential (atmospheric pressure), gravitational potential (usually insignificant in soils), and overburden potential (weight of overlying matter) make only small contributions to water available for physiological processes (Papendick and Campbell 1981). They also probably affected all hatcheries equally and can thus be considered unimportant in this context. Unfortunately, the tensiometer used in this experiment was not capable of blocking the flow of ions, so the water potential data presented here are limited to matric potential + pressure potential and do not include the osmotic potential component of total water potential. This is an important distinction and becomes a factor in highly saline sand.

Elevated salinity lowers soil osmotic potential, and therefore total water potential, in the presence of a semipermeable barrier (i.e., eggshells) (Rimkus et al. 2002). In the absence of a strong (i.e., highly negative) osmotic potential, high moisture and water potential is generally expected to allow sea turtle eggs to retain and even absorb water (Lohmann et al. 1997). Although matric potential is generally the largest component of water potential in unsaturated sand (Papendick and Campbell 1981), it is likely that the matric potential component of total water potential in the irrigated hatcheries was overwhelmed by the osmotic forces resulting from the influx of highly saline seawater used for irrigation. This may explain why many of the irrigated eggs became severely desiccated despite the high moisture and water potential recorded in the hatchery sand.

### 4.2. Biotic responses to seawater irrigation

#### 4.2.1. Embryonic development

The pattern of the response of *C. mydas* eggs and embryos to large-scale and long-term irrigation with seawater highlights the strong negative effect such an irrigation regimen has on embryonic development. Our observation that intensely irrigated clutches suffered almost complete failure aligns with the wealth of evidence indicating that seawater inundation and ground-water flooding of sea turtle nests has a negative impact on sea turtle nests and often leads to reproductive failure. However, an unexpected result was the large proportion of embryos in the irrigated hatcheries that survived to the most advanced field stage of development (stage 3). The field staging technique used in this study divides embryonic development into easily identifiable benchmarks, but the field stages differ in duration and in the number of developmental stages they encompass. Field stage 3, at which over 60% of unhatched eggs in the irrigated hatcheries died, begins at approximately 45% of incubation time and continues until hatching (Miller 1985, Rafferty et al. 2011). Field stage 3 thus encompasses a wide range of development and its temporal resolution is coarse. We observed that many of the stage 3 embryos had visible scales on the flippers and head and in some cases the embryos were larger than the yolk. These morphologies identify embryos as being aged between 60–86% of the incubation period (Miller 1985), which indicates that these embryos survived for a substantial proportion of their development in conditions of very high salinity, moisture, and water potential. This surprising result suggests an encouraging robustness to salinity in embryos of *C. mydas* that has heretofore been lacking in the published literature. There is ample evidence that irrigation with seawater does not always result in developmental failure (Limpus et al. 2020, Smith et al. 2021, Young et al. 2023), but little research has been dedicated to identifying the upper limit in terms of the volume of seawater and the frequency of irrigation that sea turtle eggs can withstand. Addressing this knowledge gap is important in order to determine the utility and parameters of a sea-water irrigation regime for conservation purposes.

### 4.3 Recommendations for researchers and managers

Although the methods employed in this study clearly surpassed the tolerance limit of *C. mydas* embryos for seawater infiltration into their nests, we think that sensible adjustments to the regimen could achieve the goal of temperature reduction without causing unacceptable developmental failure or pre-emergent hatchling mortality. One adjustment may encompass a reduction in the number of times irrigation is applied, with fewer applications over a briefer time period, for example, so that nests which happen to be at the correct age at that time would be irrigated during the thermosensitive period. Another option, if fresh water is available, would intersperse seawater irrigation with fresh water irrigation. If fresh water were occasionally applied, either on the final watering or alternating with irrigation with seawater, it would accelerate the natural process by which the beach sand returns to its baseline salinity, hopefully reducing stress on the embryos. It is likely that an irrigation regimen that integrates a combination of these adjustments may prove to be the most effective approach to minimise embryonic mortality. If irrigation with seawater is to be of continuing interest to conservation, further research into the use of these protocols and how to best reduce embryonic mortality is essential. In that way, we think that large-scale seawater irrigation may yet prove to be a viable management intervention for management of climate change impacts on sea turtle populations.

## 5. Conclusions

Managing sea turtle populations in the face of increasing global temperature may require the implementation of interventions aimed at reducing heat-induced embryonic mortality and over-feminisation of populations (Pike 2014, Hays et al. 2017, Laloë et al. 2017). We evaluated the efficacy of two large-scale and prolonged regimens of irrigation using seawater to reduce sand temperature in clutches of green turtle eggs and the impacts of those regimens on nest environmental conditions, embryonic development, and developmental success. The more intense irrigation regimen effectively reduced the temperature of clutches by an average of 1.3 °C and both intense and moderate irrigation frequently resulted in over 3 °C of cooling. Sand moisture, water potential, salinity, and clutch oxygen concentration were significantly higher in irrigated than unirrigated hatcheries, but clutch oxygen availability appeared to be unaffected by irrigation. Hatching success in the control hatchery was similar to natural nests at the study site but was very low in both irrigated groups. Of particular interest is the discovery that most of the embryos experienced mortality at later stages of development. This indicates that minor alterations to the irrigation protocol used in this study may allow irrigation with seawater to achieve the desired temperature change without unacceptably high embryonic mortality and provide evidence of its potential as a viable and effective management intervention for sea turtle conservation. Some suggestions for modifications to the protocols used in this study include fewer bouts of irrigation and alternating between fresh water and seawater when practicable. However, the full effects of such interventions, such as how changes to the nesting habitat will impact the ability of sea turtles to adapt to climate change, remain uncertain (Fuentes et al. 2023). Future research is essential in order to determine the most effective regimen of irrigation involving seawater and the extent to which sea turtle eggs and embryos can withstand irrigation with seawater.

## Acknowledgements

Funding for this project was provided by the Holsworth Wildlife Research Endowment, the Ecological Society of Australia, and Monash University. This work was part of David M. Adams’ PhD thesis, during which he was financially supported by an Australian Government Research Training Program (RTP) scholarship. We acknowledge the Gooreng Gooreng, Gurang, Bailai and Taribelang Bunda peoples as traditional custodians of Heron Island and the surrounding waters. We are grateful to the staff at the Heron Island Research Station for their assistance and support during our visits to the island. We thank the sea turtle monitoring volunteers of the Queensland Department of Environment, Tourism, Science, and Innovation for their assistance during the nesting season, and to Briana Gibbs for assistance with hatchery construction and egg collection.

---

**Supplementary Figure 1.**
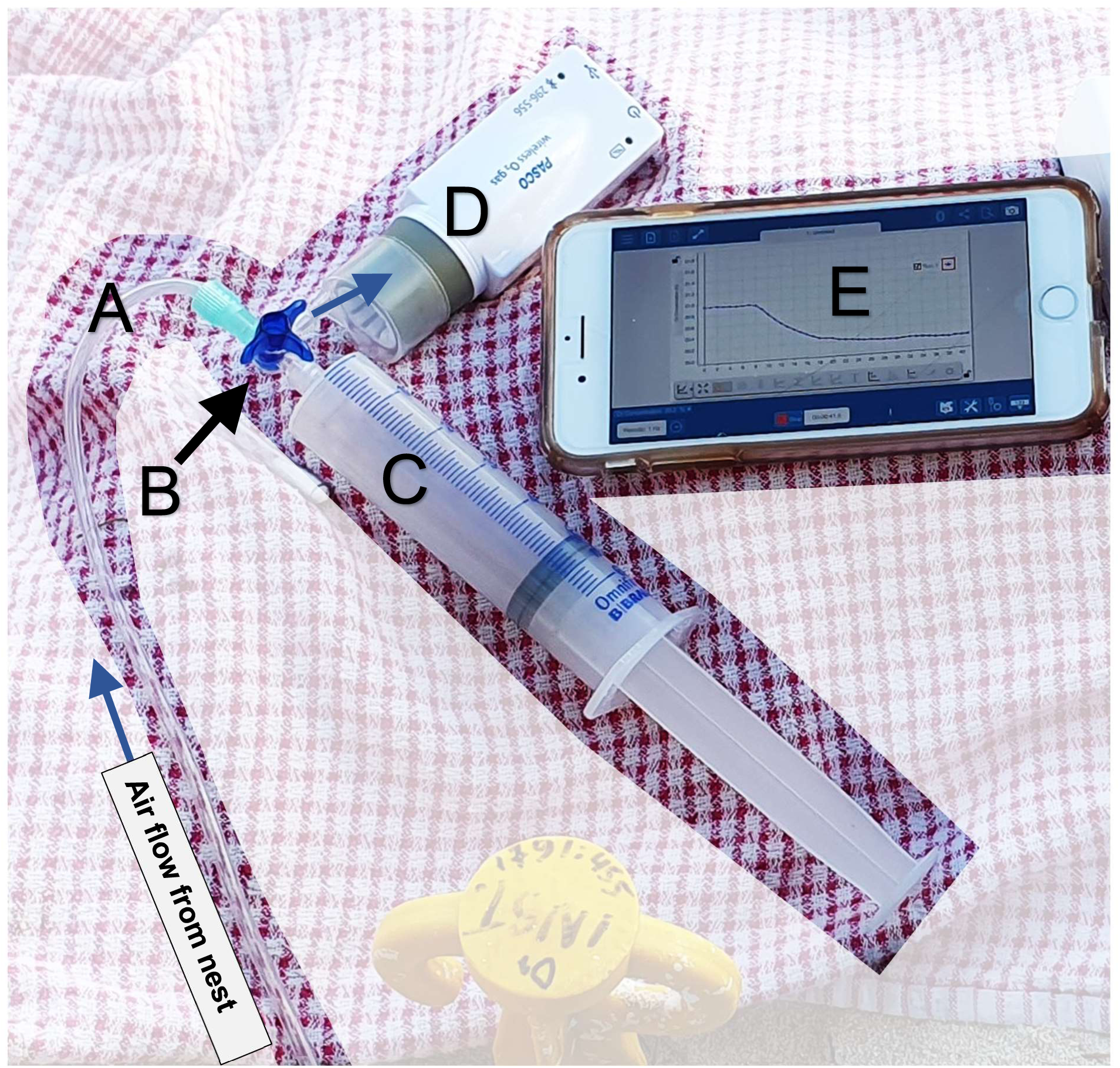
Clutch oxygen sampling equipment. Air was drawn from the nest through the sampling tube (A) (3 mm lumen) through the 3-way stopcock (B) into the syringe (C). Stopcock was then turned to allow air to flow out through the oxygen detector (D). In this image the syringe is filled with 50 ml of nest air, stopcock is shown aligned to inject the sample of air through the sensor. Blue arrows indicate the direction of one-way air flow. Data from the oxygen sensor is recorded on the tablet (E) for later analysis.

**Supplementary Table 1.** Post hoc pairwise comparisons of oxygen content between clutches of green turtle (*Chelonia mydas*) eggs irrigated with seawater. Shown here are *p*-values of post hoc comparisons of daily oxygen concentration between clutches subjected to the same irrigation regimen. *P*-values in the table were calculated using Bonferroni-corrected pairwise Dunn’s tests, which were performed after significant results from Kruskal-Wallis rank sum tests. Results of Kruskal-Wallis tests are listed at the top of each section. *P*-values > 0.05 are not considered statistically significant (ns), significant *p*-values ≤ 0.05 are listed in the table.

| **Control (unirrigated) Clutches** ($\chi^2 = 22.7$, df = 7, $p = 0.002$) | | | | | | | | |
| --- | --- | --- | --- | --- | --- | --- | --- | --- |
|  | Nest 1 | Nest 2 | Nest 3 | Nest 4 | Nest 5 | Nest 6 | Nest 7 | Nest 8 |
| Nest 2 | ns |  |  |  |  |  |  |  |
| Nest 3 | ns | ns |  |  |  |  |  |  |
| Nest 4 | ns | ns | ns |  |  |  |  |  |
| Nest 5 | ns | ns | ns | ns |  |  |  |  |
| Nest 6 | ns | ns | ns | ns | ns |  |  |  |
| Nest 7 | ns | ns | ns | ns | ns | ns |  |  |
| Nest 8 | < 0.01 | 0.01 | ns | ns | ns | ns | ns |  |

| **Intense Irrigated Clutches** ($\chi^2 = 115$, df = 8, $p < 0.001$) | | | | | | | | |
| --- | --- | --- | --- | --- | --- | --- | --- | --- |
|  | Nest 1 | Nest 2 | Nest 3 | Nest 4 | Nest 5 | Nest 6 | Nest 7 | Nest 8 |
| Nest 2 | ns |  |  |  |  |  |  |  |
| Nest 3 | < 0.01 | ns |  |  |  |  |  |  |
| Nest 4 | < 0.001 | 0.03 | ns |  |  |  |  |  |
| Nest 5 | ns | ns | ns | < 0.001 |  |  |  |  |
| Nest 6 | ns | < 0.01 | < 0.001 | < 0.001 | ns |  |  |  |
| Nest 7 | ns | < 0.001 | < 0.001 | < 0.001 | < 0.01 | ns |  |  |
| Nest 8 | ns | ns | < 0.01 | < 0.001 | ns | ns | ns |  |
| Nest 9 | ns | ns | ns | 0.01 | ns | 0.02 | < 0.001 | ns |

| **Moderate Irrigated Clutches** ($\chi^2 = 142$, df = 8, $p < 0.001$) | | | | | | | | |
| --- | --- | --- | --- | --- | --- | --- | --- | --- |
|  | Nest 1 | Nest 2 | Nest 3 | Nest 4 | Nest 5 | Nest 6 | Nest 7 | Nest 8 |
| Nest 2 | ns |  |  |  |  |  |  |  |
| Nest 3 | ns | ns |  |  |  |  |  |  |
| Nest 4 | 0.02 | ns | 0.03 |  |  |  |  |  |
| Nest 5 | < 0.001 | < 0.001 | < 0.001 | ns |  |  |  |  |
| Nest 6 | ns | ns | ns | ns | ns |  |  |  |
| Nest 7 | < 0.001 | < 0.001 | < 0.001 | < 0.001 | ns | < 0.001 |  |  |
| Nest 8 | < 0.001 | < 0.001 | < 0.001 | ns | ns | ns | 0.04 |  |
| Nest 9 | < 0.001 | < 0.001 | < 0.001 | 0.02 | ns | < 0.001 | ns | ns |

**Supplementary Figure 2.**
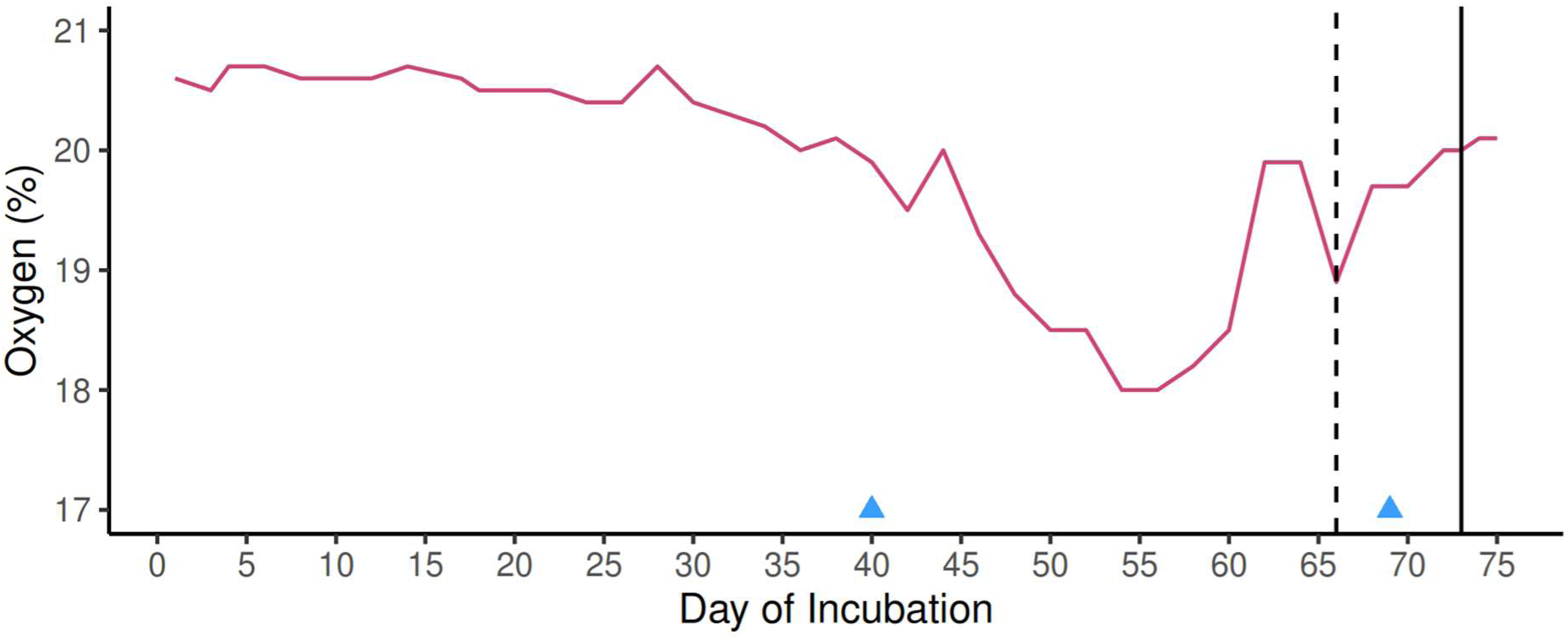
Example pattern of daily oxygen content in unirrigated nests. Shown here is nest 6, a relocated clutch of 83 eggs. Hatching success of this nest was 90.4%, emergence success was 88.0%. Dashed line indicates estimated day of hatching (day 66), solid line indicates day of emergence (day 73). Inverted triangles indicate significant rainfall events (> 15 mm). Note decline in oxygen content beginning approximately halfway through incubation and period of quiescence, indicated by a rise in oxygen, in the days before hatching. Hatching was estimated to occur after this period of quiescence and is indicated by the final sharp drop in oxygen content, which occurred ∼7 days before emergence.

